# Targeting Sec61 to Silence Pathogenic Light-Chain Production in AL Amyloidosis

**DOI:** 10.64898/2026.09.10.750585

**Authors:** Daniela Ricci, Chloé Veront, Irene Valcozzena, Merlin Després, Alice Nevone, Marine Oberkampf, Mariette Matondo, Karen Druart, Thibaut Douché, Sarra Loulizi, David Hardy, Florence Bugault, Manon Capdeville, Mario Nuvolone, Caroline Demangel

## Abstract

Unstable immunoglobulin light chains (LCs) secreted by clonal plasma cells drive organ damage in AL amyloidosis, yet no therapy exists to directly suppress their biosynthesis. Here, we targeted the Sec61 translocon to block LC biogenesis. In patient-derived plasma cells, the Sec61 inhibitor mycolactone redirected nascent LCs to proteasomal degradation, inducing proteotoxic stress that synergized with the proteasome inhibitor bortezomib. *In vivo*, this combination reduced plasma-cell engraftment and LC secretion but caused systemic toxicity, underscoring the need for Sec61 inhibitors with client selectivity, i.e. the ability to block only the translocation of a restricted number of Sec61 substrates. Proteomic profiling of a model patient-derived plasma cell line treated with the synthetic Sec61 inhibitors A317 and A347 revealed intermediate- and narrow-spectrum client selectivity, respectively, while retaining activity against the pathogenic LC. However, unlike A317, the more selective A347 failed to inhibit diverse amyloidogenic LCs in a signal-peptide reporter assay. In primary plasma cells from a cohort of patients with AL amyloidosis, A317 consistently reduced pathogenic LC production while sparing matched non-malignant bone marrow cells. Together, these findings establish Sec61 as a therapeutic target in AL amyloidosis and highlight the need to balance client selectivity with coverage of diverse amyloidogenic LC signal peptides.

## INTRODUCTION

Systemic immunoglobulin light-chain (AL) amyloidosis is a protein misfolding disorder caused by clonal plasma cells producing unstable monoclonal immunoglobulin light chains (LCs), with an incidence of 12 cases per million individuals (Bianchi et al., 2026). Misfolded LCs assemble into amyloid fibrils that deposit in vital organs, including the heart, kidneys, liver, and peripheral nervous system, causing progressive dysfunction and high mortality. Despite major advances in plasma cell-directed therapies, including proteasome inhibitors, immunomodulatory drugs, and monoclonal antibody-based regimens, many patients fail to achieve durable hematologic responses, and organ recovery remains limited. Approaches that directly target the production of amyloidogenic LCs are still lacking.

LCs enter the endoplasmic reticulum (ER) through co-translational translocation mediated by the Sec61 translocon, the central gateway of the secretory pathway (Itskanov and Park, 2023; Sundaram et al., 2025). Because ER entry is an essential early step in LC maturation and secretion, disrupting this process offers a direct strategy to reduce the generation of aggregation-prone precursors. Pharmacological Sec61 inhibition is emerging as a means to modulate secretory protein biogenesis, as substrates that fail to translocate are redirected to cytosolic proteasomal degradation after translation termination (Baron et al., 2016; Grotzke et al., 2017; Guenin-Mace et al., 2015; Ricci and Demangel, 2023). This approach is particularly attractive in AL amyloidosis, where disease progression relies on continuous secretion of amyloidogenic LC by clonal/tumoral plasma cells.

We previously showed that mycolactone, a Sec61 blocker, induces lethal proteotoxic stress in multiple myeloma (MM) cells and synergizes with proteasome inhibition (Domenger et al., 2023, 2022). However, by broadly inhibiting Sec61-dependent translocation, mycolactone disrupts proteostasis and compromises viability across cell types, limiting its therapeutic window (Morel et al., 2018). In contrast, other described Sec61 inhibitors preferentially impair the biogenesis of specific client proteins, thus causing less extensive proteostasis disruption and reduced cytotoxicity (Garrison et al., 2005; Rehan et al., 2023; Wenzell et al., 2024). These findings suggest that controlled Sec61 inhibition may enable targeted degradation of disease-relevant substrates. A317 and A347 are synthetic Sec61 inhibitors developed by Kezar Life Sciences to primarily target PD-1, with varying degrees of cross-reactivity with the other Sec61 clients HER3, TNFα and IL-2 (WO2020176863A1). However, their proteomic signature across the human Sec61 clientome and their therapeutic potential remain unknown.

In this study, we evaluated mycolactone together with A317 and A347 in cellular, animal, and patient-derived models of AL amyloidosis to test whether client-selective inhibition of LC translocation could reduce pathogenic LC production with limited toxicity. Our findings demonstrate that client-selective Sec61 inhibition can effectively suppress pathogenic LC production while improving tolerability compared with broad Sec61 blockade by mycolactone. They also identify germline-specific LC signal peptide diversity as an important determinant of therapeutic response.

## RESULTS

### Sec61 inhibition impairs LC secretion and viability of AL cells *in vitro* and *in vivo*

To assess whether Sec61 inhibition can suppress the production of pathogenic LCs, we used the AL patient-derived ALMC-1 plasma cell line, which secretes amyloidogenic λ LCs both as free LCs and as part of an intact monoclonal antibody (Arendt et al., 2008). Cell treatment with mycolactone (Myco; 0.8-100 nM) for 24 h reduced LC secretion in a dose-dependent manner (IC₅₀ = 33 nM) and markedly decreased cell viability (Fig. 1A). Because plasma cells are highly dependent on proteostasis, we next examined whether Sec61 blockade could enhance the activity of the proteasome inhibitor bortezomib (BZ). BZ alone (0.4-50 nM) also strongly reduced both LC secretion (IC₅₀ = 6 nM) and cell viability in ALMC-1 cells (Fig. 1A). Combining BZ with 25 nM mycolactone produced even greater reductions in both LC secretion and viability (Fig. 1B), and drug-interaction analysis using the Highest Single Agent (HSA) model confirmed synergy between Myco and BZ across a broad range of compound concentrations (Supplementary Fig. 1A). Together, these results indicate that Sec61 blockade inhibits LC translocation, thereby preventing LC biogenesis, and synergizes with proteasome inhibition to enhance killing of AL plasma cells.

**Figure 1.**
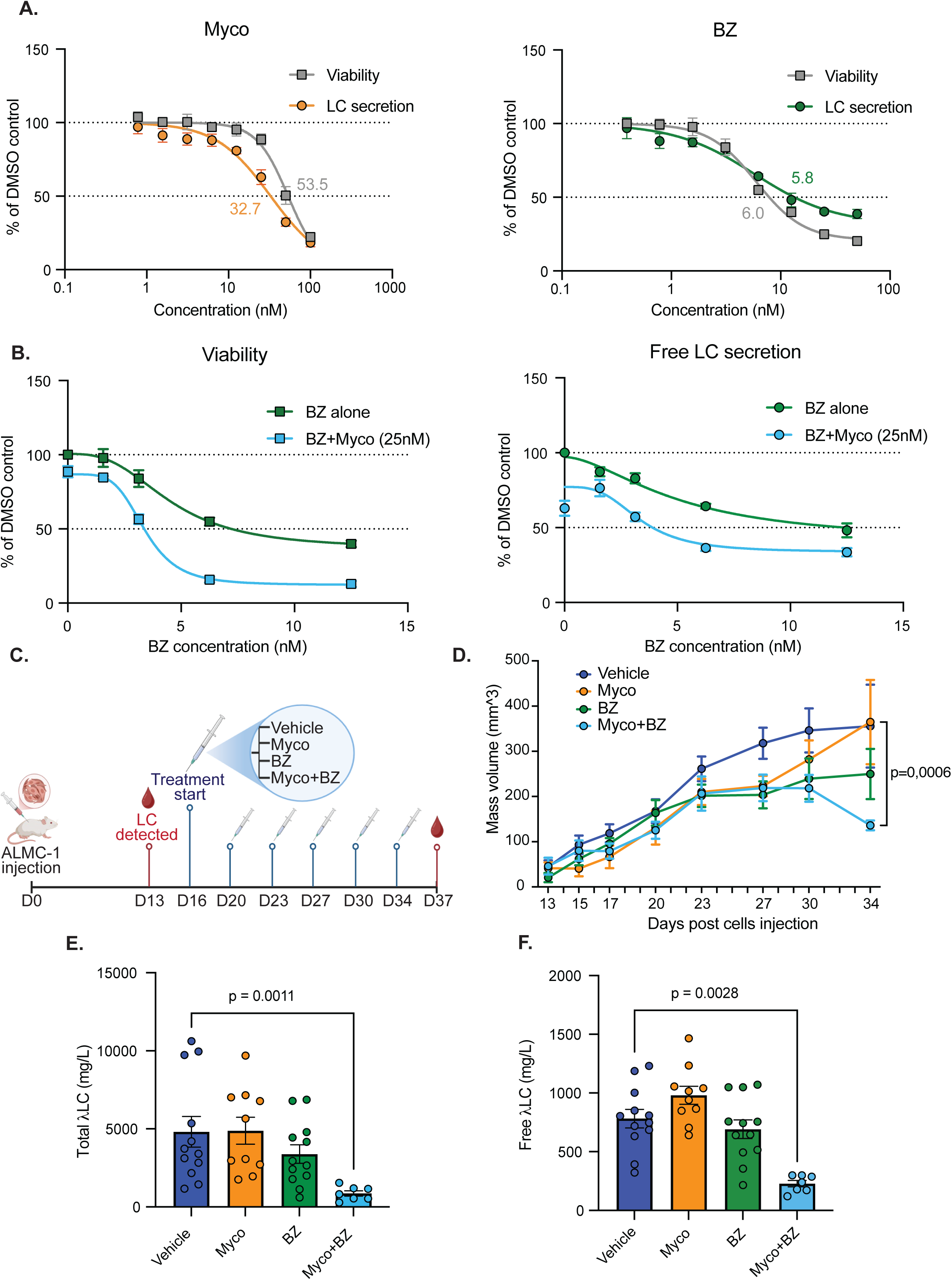
Sec61 inhibition reduces LC secretion and compromises AL plasma-cell viability in vitro and in vivo. (A-B) Dose-dependent effects of mycolactone (Myco) and bortezomib (BZ) alone **(A)** or in combination **(B)** on ALMC-1 cell viability and free LC secretion after 24 h. Data are mean ± SD from three independent experiments. **(C)** ALMC-1 xenograft study design. NSG mice bearing established ALMC-1 grafts were randomized on day 16 to vehicle (n = 12), BZ (0.5 mg/kg; n = 12), Myco (0.3 mg/kg; n = 11), or Myco plus BZ (n = 11), administered intraperitoneally twice weekly. **(D)** ALMC-1 graft growth during treatment. Data are mean ± SEM; two-way ANOVA with Tukey’s multiple-comparisons test. **(E–F)** Total **(E)** and free **(F)** serum LC concentrations on day 37. Bars represent mean ± SEM; Kruskal-Wallis test with Dunn’s multiple-comparisons test.

We next compared mono-versus combination therapy *in vivo*. ALMC-1 cells mixed with Matrigel were implanted subcutaneously into NOD/SCID/IL2rγnull (NSG) mice to generate xenografts and assess plasma cell engraftment and therapeutic response (Fraser et al., 2022). Graft volume was measured every three days and circulating pathogenic LC levels were quantified by ELISA. Circulating LCs became detectable in all mice by day 13. On day 16, mice were randomized to receive vehicle, Myco (0.3 mg/kg), BZ (0.5 mg/kg), or the combination. Treatments were administered intraperitoneally twice weekly (Fig. 1C). Combination therapy reduced plasma cell engraftment from day 30 onward (Fig. 1D) and significantly decreased both total and free serum LC concentrations (Fig. 1E, F). In contrast, neither single-agent treatment significantly affected graft growth or circulating LC concentrations under the conditions tested (Fig. 1D-F). Although continuous weight monitoring and histopathological examination of the heart, kidneys, liver, and pancreas revealed no evidence of treatment-related toxicity, the Myco+BZ combination treatment resulted in the death of 4 of 11 animals (Table I, Supplementary Fig. 1B-D). However, consistent with reduction in engraftment (Fig. 1D), histological images of excised tumor masses showed altered tissue architecture in the combination-treated group compared with the single-agent and vehicle groups, suggestive of selective cytotoxicity toward plasma cells within the graft (Supplementary Fig. 1E). Therefore, despite its potent anti-LC activity *in vivo*, combined Sec61 inhibition and BZ treatment appear to have a limited therapeutic window, prompting us to evaluate client-selective Sec61 inhibitors.

**Table I.** Histopathology summary.

| Treatment | n | Heart | Kidney | Liver | Pancreas / overall interpretation |
| --- | --- | --- | --- | --- | --- |
| **Vehicle** | 12 | 11 normal; 1 minimal interstitial oedema and mild hyperaemia, without inflammatory or degenerative lesion. | Normal in all examined mice. | Normal in all examined mice. | Normal in all examined mice. No systemic treatment-related lesions reported. |
| **Mycolactone** | 11 | Normal in all examined mice. | Normal in all examined mice. | Normal in all examined mice. | Normal in all examined mice. No treatment-related organ toxicity reported. |
| **Bortezomib** | 12 | 10 normal; 1 focal fibrino-haemorrhagic intracardiac mural thrombus or adherent necrotic debris; 1 minimal reactive oedema/disorganisation, without myocarditis or necrosis. | Normal in all examined mice. | Normal in all examined mice. | Normal in all examined mice. Isolated cardiac observations only; no consistent treatment-related pattern reported. |
| **Mycolactone + bortezomib** | 7 | Normal in all examined mice. | Normal in all examined mice. | Normal in all examined mice. | Normal in 6/7 mice; pancreas not sampled in 1 mouse. No treatment-related organ toxicity reported in evaluable tissues. |

### Client-selective Sec61 inhibitors suppress pathogenic LC secretion with reduced cytotoxicity

A317 and A347 are chemically related synthetic Sec61 inhibitors initially developed to prevent the biogenesis of the T-cell checkpoint receptor PD-1. We assessed whether these compounds also affect Sec61-dependent production of the pathogenic LC secreted by ALMC-1 cells. Although less potent than mycolactone (Fig. 1A), A317 and A347 reduced LC secretion in a dose-dependent manner, with IC₅₀ values of 0.1 µM and 1.1 µM, respectively (Fig. 2A). In contrast to mycolactone, A317, and particularly A347, substantially inhibited LC secretion at concentrations that did not impair cell viability (Fig. 2A).

**Figure 2.**
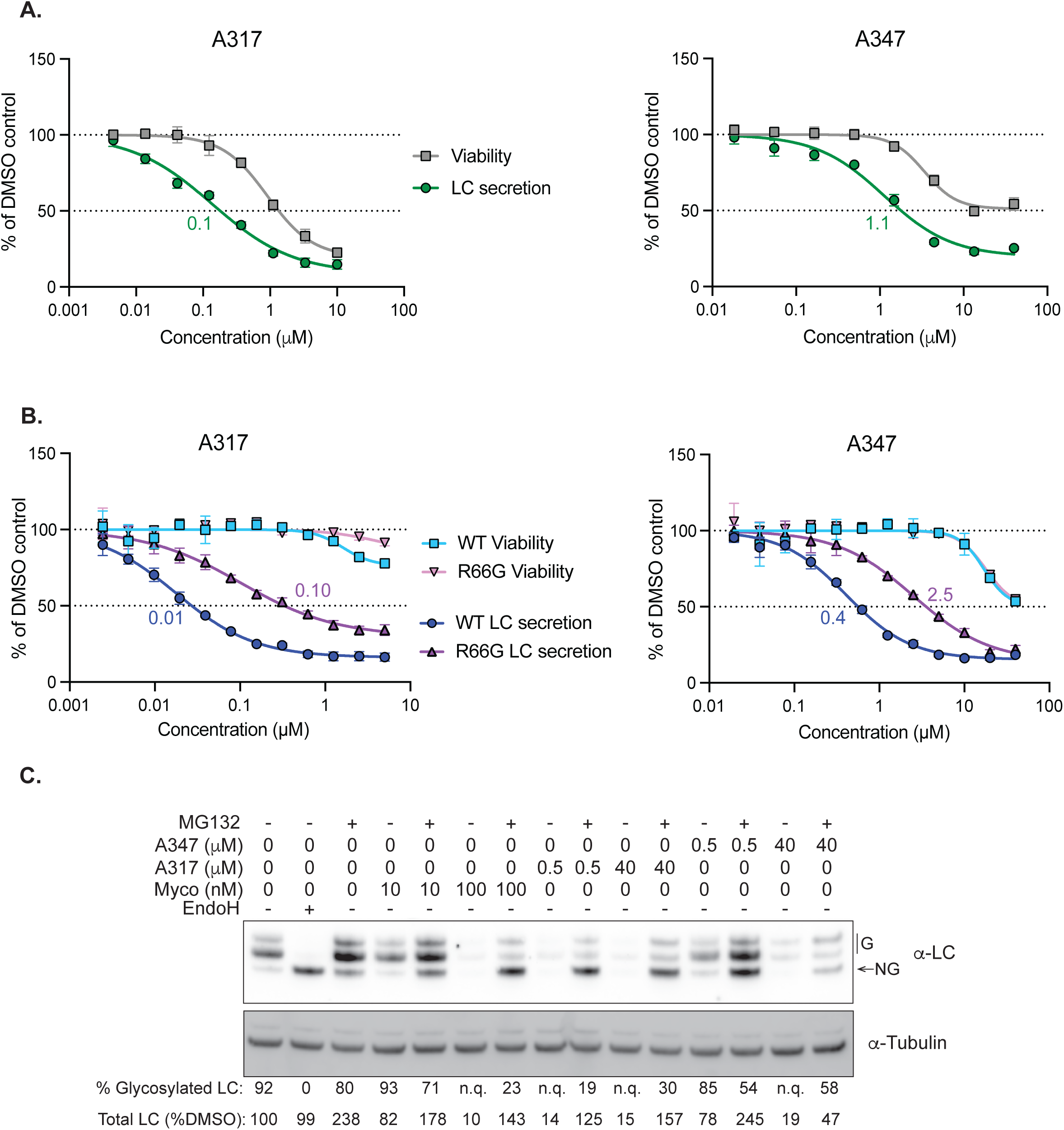
Client-selective Sec61 inhibitors suppress pathogenic LC secretion with reduced cytotoxicity. **(A)** Dose-dependent effects of A317 and A347 on ALMC-1 cell viability and free LC secretion after 24 h. Data are mean ± SD from three independent experiments, with best-fit IC₅₀ values. **(B)** Dose-dependent effects of A317 and A347 on cell viability and free LC secretion by Flp-In T-REx 293 LC-NanoLuc cells expressing wild-type (WT) or R66G Sec61 after 24h. Data are mean ± SD from two independent experiments; relative to vehicle controls, with best-fit IC₅₀ values. **(C)** HEK293 LC-opsin cells were treated for 6 h with mycolactone (Myco), A317 and A347, with or without MG132. EndoH digestion identifies glycosylated, ER-translocated LC species. % glycosylated LC = glycosylated LC / (glycosylated + non-glycosylated LC); total LC was normalized to α-tubulin and expressed relative to DMSO. n.q., not quantifiable; G: glycosylated; NG: not glycosylated. Immunoblots are representative of two independent experiments.

To confirm that this effect resulted from Sec61 inhibition, we generated HEK293 cell lines expressing the ALMC-1-derived pathogenic LC together with either wild-type Sec61 (Sec61-WT) or the inhibitor-resistant Sec61-R66G mutant. This mutation has been reported to prevent binding of all currently characterized Sec61 inhibitors (Itskanov et al., 2023; Luesch and Paavilainen, 2020). Consistent with previous findings, expression of Sec61-R66G markedly reduced mycolactone-mediated inhibition of LC secretion relative to Sec61-WT controls (Supplementary Fig. 2). Similarly, expression of Sec61-R66G significantly reduced A317- and A347-induced inhibition of LC secretion, as evidenced by a marked rightward shift in the IC₅₀, confirming that both compounds act through Sec61 engagement (Fig. 2B).

We next investigated whether A317- and A347-mediated LC suppression resulted from impaired ER translocation and subsequent degradation of nascent LC. Because the LC itself lacks endogenous N-linked glycosylation sites, we generated HEK293 cells expressing the pathogenic LC fused at its C-terminus to an opsin-derived glycosylation reporter containing two N-linked glycosylation sites, which are modified only upon ER entry, thereby providing a readout of LC translocation into the ER (LC-opsin) (Roboti et al., 2021). In untreated cells, LC-opsin was detected predominantly as a glycosylated species, with a minor non-glycosylated form migrating at the same molecular weight as LC-opsin in Endo H-treated lysates (Fig. 2C). Treatment with mycolactone (Myco; 10-100 nM, 6 h) dose-dependently reduced glycosylated LC-opsin levels. This effect was rescued by the proteasome inhibitor MG132, showing that Sec61 blockade prevents ER entry and redirects nascent LC-opsin for proteasomal degradation (Fig. 2C). Likewise, A317 and A347, tested at 0.5 µM and 40 µM, reduced glycosylated LC-opsin levels, confirming that both compounds impair LC biogenesis by inhibiting ER translocation (Fig. 2C). Together, these results establish Sec61-mediated ER-translocation blockade as the mechanism underlying LC suppression by A317 and A347.

### Global proteomic analysis defines the Sec61 clientome targeted by A317 and A347 in ALMC-1 cells

To assess the selectivity of A317 and A347 across the Sec61 clientome in ALMC-1 cells, we performed global proteomic profiling after 24 h of treatment at IC₅₀ concentrations, conditions chosen because they produced comparable suppression of LC production while limiting secondary effects associated with reduced cell viability (Fig. 2A). Mycolactone was excluded, as its cytotoxicity could not be dissociated from its broad Sec61 inhibitory activity (Fig. 1A). ALMC-1 cells were treated with A317 or A347 in five technical replicates, and only proteins consistently and significantly modulated across replicates were retained for analysis. A317 altered more proteins than A347 (146 versus 82 significantly modulated proteins, respectively), including 52 versus 29 downregulated Sec61 clients (Fig. 3A-B), and upregulated more proteins overall (41 versus 26). Despite limited overlap at the individual-protein level, proteins modulated by both compounds converged on a shared proteomic footprint, differing primarily in the magnitude of the response (Fig. 3C).

**Figure 3.**
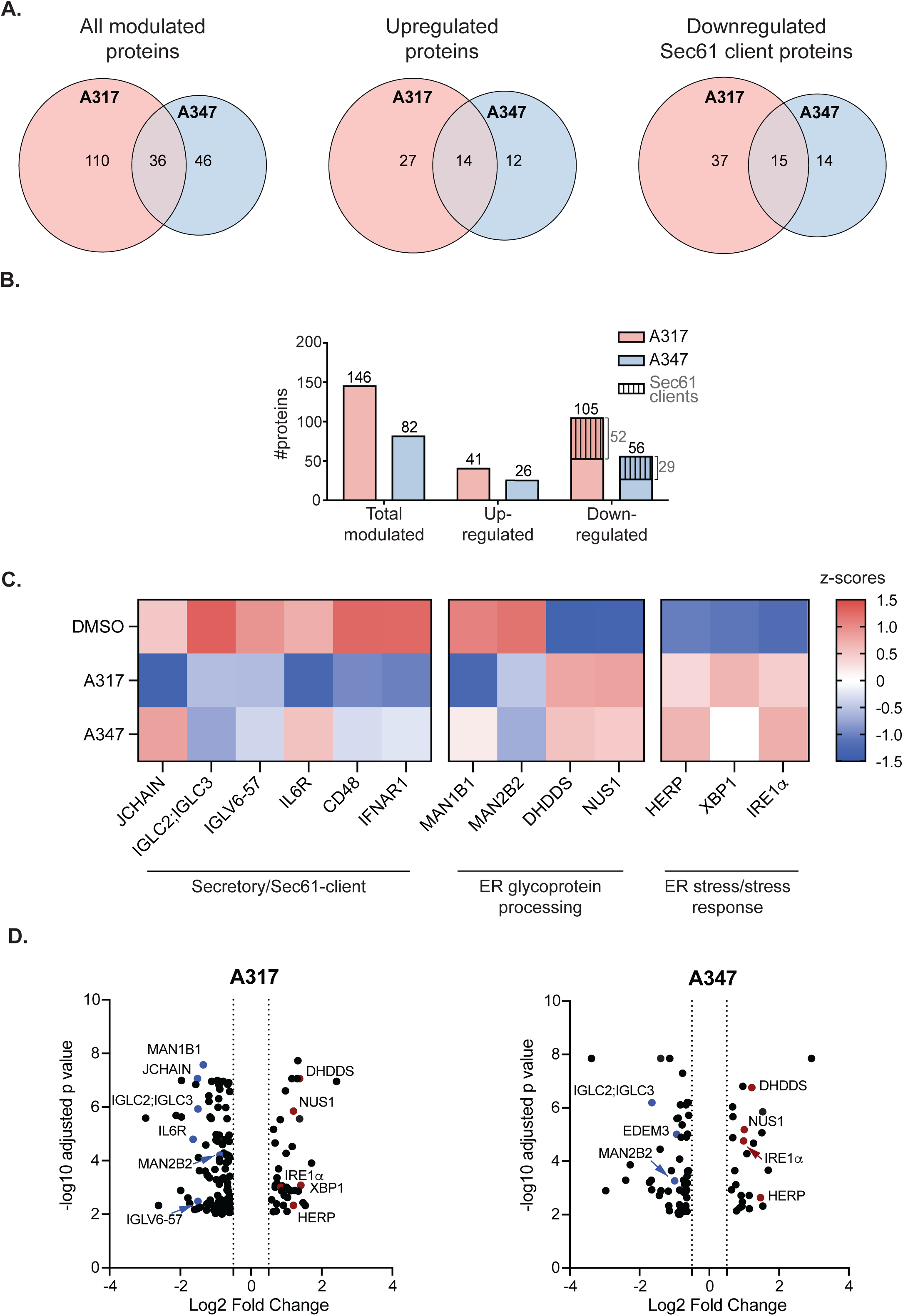
Global proteomic analysis defines the Sec61 clientome targeted by A317 and A347. **(A)** Overlap between proteins significantly modulated by A317 and A347, shown separately for all regulated proteins, upregulated proteins, and downregulated Sec61 clients. **(B)** Number of significantly modulated proteins per treatment; hatched regions indicate downregulated Sec61 clients. **(C)** Heatmap of selected proteins involved in ER stress/stress-response pathways, ER glycoprotein processing, and secretory/Sec61-client biology. Color reflects relative abundance (row z-scores of log2-transformed values) across DMSO, A317, and A347-treated samples. **(D)** Volcano plots of significant protein-abundance changes with A317 or A347 relative to DMSO. Selected ER stress, proteostasis, and Sec61-client proteins are labelled; downregulated and upregulated proteins are shown in red and blue, respectively.

In line with our cellular assays, both compounds significantly reduced LC biogenesis, as reflected by decreased abundance of immunoglobulin LC constant-region peptides (IGLC2; IGLC3). A317 more broadly affected immunoglobulin and additional Sec61-client components, including the IgA monomer linker JCHAIN, peptides from the variable region of the ALMC-1-derived immunoglobulin IGLV6-57, and the plasma cell pro-survival receptor IL6R, while A347 showed weaker or absent effects on several of these targets (Fig. 3C, D). Both inhibitors also reduced the lysosomal α-mannosidase MAN2B2, whereas A317 additionally reduced the ER α-mannosidase MAN1B1, suggesting common but quantitatively distinct alterations in mannose trimming of glycoprotein-derived N-glycans. Other Sec61-client proteins, such as CD48 and IFNAR1, also showed stronger reduction with A317 than with A347 (Fig. 3C), consistent with the broader client spectrum of A317.

Both compounds upregulated the stress-induced ERAD component HERP, and the ER stress sensor IRE1α, with A317 additionally increasing XBP1. These changes indicate activation of the unfolded protein response (UPR), as previously observed in mycolactone-treated cells (Morel et al., 2018). Interestingly, DHDDS and NUS1, which form the cis-prenyltransferase complex required for dolichol synthesis, were upregulated by both compounds, potentially reflecting a compensatory response to perturbed dolichol-dependent N-glycosylation. Consistent with this interpretation, downregulation of EDEM3, MAN1B1 and MAN2B2 further pointed to remodeling of ER glycoprotein processing pathways. Overall, this analysis identified A317 and A347 as Sec61 inhibitors with intermediate and high client selectivity, respectively, with partially overlapping yet distinct proteomic signatures. The higher selectivity of A347 correlated with lower induction of stress responses and reduced proteostasis alterations.

### SP sequence determines susceptibility of amyloidogenic LCs to Sec61 inhibitors

Having established that Sec61 inhibition suppresses the secretion of amyloidogenic LC by ALMC-1 cells, we next asked whether this activity extends across the sequence diversity of LCs found in AL amyloidosis. Because Sec61 inhibitors are thought to compete with nascent signal peptides (SPs) for engagement with the Sec61 translocon during protein translocation (Rehan et al., 2023; Wenzell et al., 2024), we hypothesized that LC susceptibility would be determined by SP sequence.

We first determined whether the susceptibility of the ALMC-1 LC (IGLV6-57) could be recapitulated by its SP alone. We compared the full-length (FL) LC with an SP-only reporter comprising the N-terminal SP through the signal peptidase cleavage site, followed by six residues of the native mature sequence. Each construct was fused to NanoLuciferase (nLuc) and expressed in Flp-In™ T-REx™ 293 cells (Fig. 4A). GFP was co-expressed from the same transcript via an internal ribosome entry site (IRES) as an internal control. Following Sec61-dependent ER translocation and SP cleavage, nLuc is secreted into the culture supernatant, providing a quantitative readout of translocation efficiency and SP-specific inhibitor sensitivity. Reporter cells were treated with increasing concentrations of mycolactone, A317, or A347 for 24 h, and IC₅₀ values were calculated relative to untreated controls (Fig. 4B). Mycolactone produced superimposable dose-response curves for the SP-only and FL constructs, confirming that susceptibility of LC biogenesis to Sec61 inhibition and its cytotoxic consequences are fully encoded by the SP. Likewise, the SP reporter closely recapitulated the inhibitory profiles of A317 and A347 observed with the FL LC, validating this simplified system for assessing SP-specific susceptibility.

**Figure 4.**
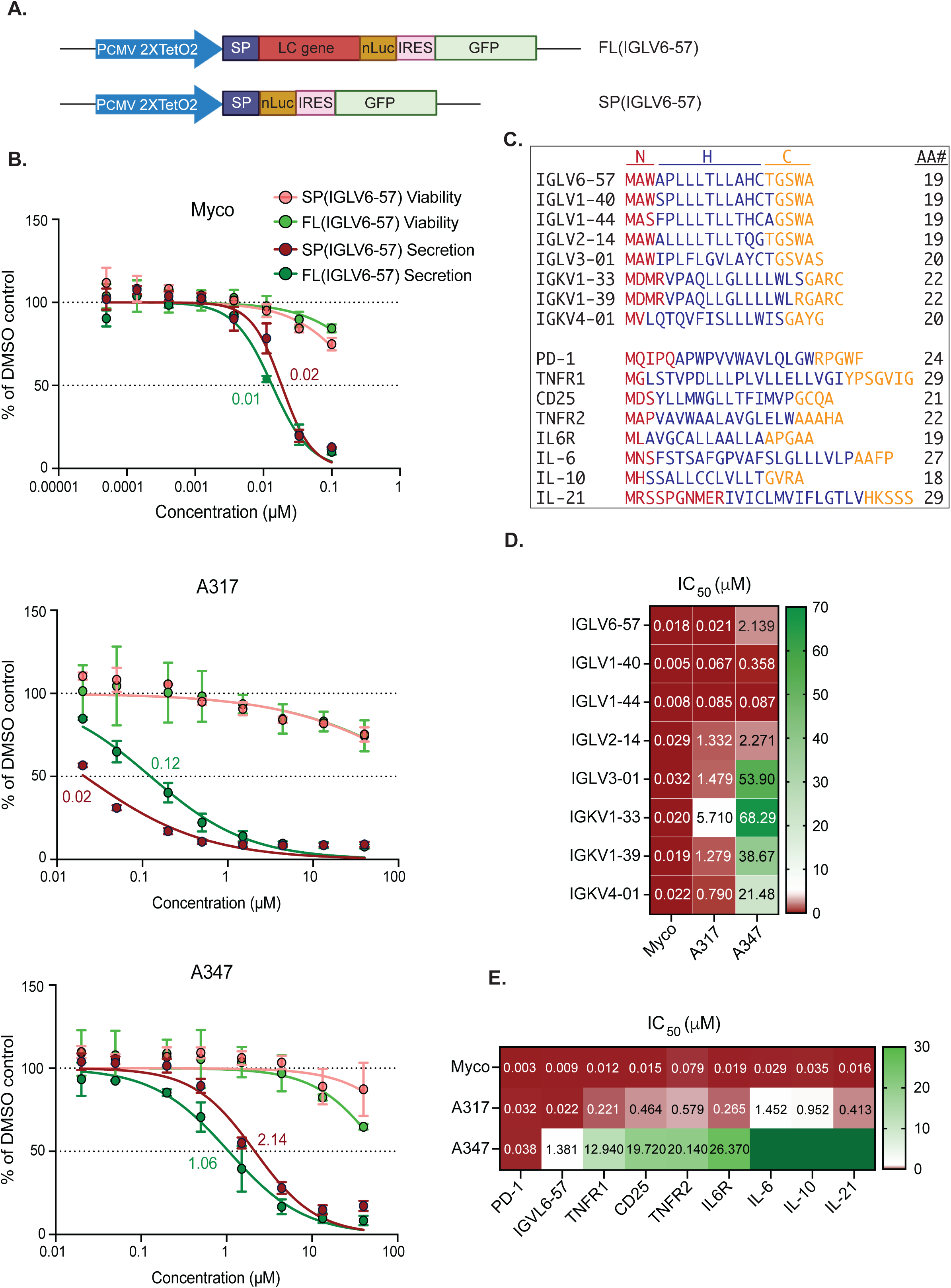
SP sequence determines susceptibility of amyloidogenic LCs to Sec61 inhibitors. **(A)** Schematic of full-length (FL) IGLV6-57 and SP-only NanoLuc (nLuc) reporter constructs. **(B)** Comparable inhibition of FL and SP-only IGLV6-57 reporters by mycolactone (Myco), A317, and A347. Data are mean ± SD from three independent experiments; with best-fit IC₅₀ values. **(C)** Alignment of amyloidogenic LC variable-region SPs. N-, hydrophobic (H)-, and C-regions are shown in red, blue, and yellow, respectively. **(D)** IC₅₀ heatmap for mycolactone (Myco), A317, and A347 across the LC SP reporter library. n = 3 for IGLV6-57 and n = 1 for other SP reporters. **(E)** IC₅₀ heatmap for mycolactone (Myco), A317, and A347 against SP reporters from IGLV6-57, PD-1, and selected plasma-cell- and inflammation-relevant proteins. White indicates an IC₅₀ of 1 µM. Dark-green cells without numerical values indicate that inhibition was not measurable. n = 2 for PD-1, n = 3 for IGLV6-57, and n = 1 for other reporters.

We next applied this assay to a panel of amyloidogenic κ and λ LC SPs representing the most common germline origins in AL amyloidosis (Morgan et al., 2025). Although all SPs retained the canonical tripartite N/H/C architecture, they displayed substantial sequence diversity (Fig. 4C). Each SP was evaluated for susceptibility to mycolactone, A317, and A347 using the same reporter system. The resulting IC₅₀ values are summarized in the Fig. 4D heatmap. While all SPs were sensitive to mycolactone (IC₅₀, 5-32 nM), susceptibility to A317 varied considerably across sequences. It was even more heterogeneous for A347, with all κ-chain and several λ-chain SPs showing little or no measurable inhibition. These findings identify SP sequence as a major determinant of susceptibility to client-selective Sec61 inhibitors.

We next asked whether A317 and A347 similarly discriminate among other Sec61 clients relevant to plasma-cell biology and the inflammatory niche, such as TNFR1/2, CD25, IL-6 and IL-6R, IL-10 and IL-21 (Zvida-Bloch et al., 2025). PD-1, the original target used to identify A317 and A347, was included as a reference. Reporter cell lines expressing the corresponding SPs were generated and treated with increasing concentrations of mycolactone, A317, or A347 for 24 h, and IC₅₀ values for each SP-inhibitor pair are shown in the Fig. 4E heatmap. For compound-SP pairs with no measurable activity, IC₅₀ values were assigned as 1,000 µM for visualization, and are shown in dark green. As expected, mycolactone efficiently inhibited translocation of all SPs (IC₅₀<100 nM). In contrast, A317 inhibited SPs derived from plasma cell- and inflammatory niche-associated proteins at micromolar concentrations, approximately tenfold higher than those required to suppress pathogenic LC secretion or PD-1. A347 displayed a markedly narrower activity profile, producing only modest inhibition of TNFR1/2, CD25, and IL-6R and no measurable activity against the remaining SPs. Together, these findings indicate that the extent of Sec61 client selectivity shapes the therapeutic profile of Sec61 inhibitors. The intermediate selectivity of A317 preserves activity against a broad range of pathogenic LC SPs and a subset of Sec61 clients involved in plasma cell survival and inflammatory signaling, which may further contribute to therapeutic efficacy. In contrast, the greater selectivity of A347 limits off-target Sec61 inhibition but reduces activity against a substantial subset of amyloidogenic LC germline genes and disease-relevant proteins.

### Selective Sec61 inhibition prevents pathogenic light-chain biogenesis by AL patient-derived plasma cells

Data in Fig. 4 identified A317 as a favorable compromise between Sec61 client selectivity and activity against amyloidogenic LCs. To evaluate its therapeutic potential in primary patient samples, we compared A317 with mycolactone for inhibition of pathogenic LC production by AL plasma cells. Bone marrow samples from 18 patients with AL amyloidosis (including AL associated with MM) were analyzed. The cohort was predominantly λ light-chain restricted, reflecting the known enrichment of the λ isotype among patients with AL amyloidosis, and encompassed a broad range of plasma cell infiltration, circulating free LC burden, organ involvement, treatment status, and t(11;14) cytogenetic status (Table II; patients 1-18). CD138⁺ plasma cells were isolated by positive immunomagnetic selection and treated *ex vivo* with mycolactone (Myco; 25-100 nM), A317 (2.5-10 µM), or BZ (0.5 µM) for 24 h. This BZ concentration was chosen as a high *in vitro* exposure, given that BZ induces proteasome inhibition and apoptosis in MM and AL plasma-cell models at low-nanomolar concentrations under continuous exposure conditions (Oliva et al., 2017). Free LC secretion and cell viability were subsequently assessed.

**Table II.** Information about the patients included in this study. AL: AL amyloidosis; ANS: autonomous nervous system; ASCT: autologous stem cell transplantation; %BMPC: percentage of bone marrow plasma cell infiltration; BJP: Bence-Jones protein; CyBorD: cyclophosphamide, bortezomib and dexamethasone; Dara: daratumumab; dFLC difference between involved and uninvolved free light chain; EPd: elotuzumab, pomalidomide and dexamethasone; FLC: free light chain; GI: gastrointestinal; IsaDex: Isatuximab and dexamethasone; LenDex: lenalidomide and dexamethasone; MDex: melphalan and dexamethasone; MM: multiple myeloma; NA: not available; PC: plasma cells; PNS: peripheral nervous system; PR: Partial response; VGPR: very good partial response.

| Pt. | Diagnosis | Tissues/organs involved | Disease stage | Monoclonal component | H chain isotype | L chain isotype | Therapy | %CD138 in NEG1 | %CD138 in POS2 | t(11;14) | dFLC (mg/L) | %BMPC |
| --- | --- | --- | --- | --- | --- | --- | --- | --- | --- | --- | --- | --- |
| 1 | MM + AL | Soft tissues | Diagnosis | FLCλ + BJPλ | - | $\lambda$ | - | 1,25 | 85,3 | Pos | 4454, | 25 |
| 2 | AL | Cardiac | Diagnosis | IgAλ + BJPλ | A | $\lambda$ | - | 0,7 | 79,2 | Neg | 292,7 | 15 |
| 3 | MM + AL | Cardiac | Diagnosis | IgGλ + FLCλ + BJPλ | G | $\lambda$ | - | 0,084 | 85,9 | Neg | 701,83 | 20 |
| 4 | AL | Cardiac | Diagnosis | IgGλ + FLCλ + BJPλ | G | $\lambda$ | - | 0,22 | 75,3 | Pos | 113,63 | 10 |
| 5 | AL | Renal and cardiac | Diagnosis | FLCλ + BJPλ | - | $\lambda$ | - | 0,33 | 82,6 | Pos | 84,43 | 8 |
| 6 | AL | Renal | Diagnosis | IgGλ + FLCλ + BJPλ | G | $\lambda$ | - | 1,04 | > 75 | Neg | 293,44 | 10 |
| 7 | AL | Cardiac and renal | Diagnosis | IgGλ + FLCλ | G | $\lambda$ | - | 1,23 | 88 | Pos | 8,3 | 10 |
| 8 | AL | Cardiac and renal | Diagnosis | FLCλ + BJPλ | - | $\lambda$ | - | 1,59 | 88,5 | Pos | 202,13 | 5 |
| 9 | AL | Renal, soft tissues, ANS and PNS | Treated - Very Good partial response | IgGλ + FLCλ + BJPλ | G | $\lambda$ | I line:<br>Dara Len<br>Dex<br>II line:<br>EPd<br>III line:<br>ixazomib<br>Dex | 0,81 | 53,3 | Neg | 33,12 | NA |
| 10 | MM + AL | Cardiac, hepatic and renal | Diagnosis | IgGλ + FLCλ + BJPλ | G | $\lambda$ | - | 2,44 | 85,3 | Pos | 146,68 | 13 |
| 11 | AL | Cardiac | Diagnosis | FLCλ + BJPλ | - | $\lambda$ | - | 3,25 | 85 | Pos | 975,3 | 1 |
| 12 | MM +<br>AL | Cardiac | Treated -<br>Partial<br>response | FLCλ +<br>BJPλ | - | λ | I line:<br>Dara<br>LenDex;<br>DaraDex | 0,33 | 76,4 | Pos | 117,5 | NA |
| 13 | MM +<br>AL | Cardiac | Treated -<br>Partial<br>response | FLCκ +<br>BJPκ | - | κ | I line:<br>CyborD<br>II line:<br>ASCT<br>III line:<br>Dara<br>LenDex;<br>Dara Dex | 0,7 | 76,5 | Pos | 80,82 | NA |
| 14 | MM +<br>AL | Cardiac | Diagnosis | FLCλ +<br>BJPλ | - | λ | - | 2,06 | 86,9 | Pos | 277,94 | 8 |
| 15 | AL | Cardiac | Diagnosis | IgGλ<br>+ BJPλ | G | λ | - | 2,2 | 87,5 | Pos | 176,02 | 1 |
| 16 | AL | Cardiac | Diagnosis | BJPλ | - | λ | - | 0,059 | 92,1 | Pos | 143,2 | 8 |
| 17 | AL | Cardiac | Diagnosis | IgGλ +<br>FLCλ +<br>BJPλ | G | λ | - | 1,6 | 89,5 | Neg | 842,95 | 12 |
| 18 | AL | Cardiac, GI and<br>ANS | Diagnosis | IgGκ +<br>BJPκ | G | κ | - | 0,17 | 77,7 | Neg | 248,83 | 20 |
| 19 | MM +<br>AL | Cardiac and<br>renal | Diagnosis | IgAλ +<br>FLCλ +<br>BJPλ | A | λ | - | 0,59 | 92,9 | Neg | 183,2 | 25 |
| 20 | AL | Cardiac and GI | Diagnosis | FLCκ +<br>BJPκ | - | κ | - | 0,32 | 87,7 | Neg | 1134,5<br>1 | 7 |
| 21 | MM +<br>AL | Cardiac, renal<br>and PNS | Treated -<br>Progressio<br>n | IgGλ | G | λ | I line:<br>Dara<br>CyborD<br>II line:<br>MDex | 1,15 | 91,8 | Pos | 79,63 | NA |
| 22 | AL | Cardiac and<br>ANS | Diagnosis | FLCλ +<br>BJPλ | - | λ | - | 1,1 | 85,7 | Pos | 140,43 | 4 |

Both mycolactone and A317 reduced free LC secretion by >80% at the lowest concentration tested in all patient samples, whereas BZ produced a substantially smaller effect under the conditions tested (Fig. 5A, B). Of note, Sec61 inhibition consistently suppressed LC production regardless of LC isotype, disease stage, or clinical severity. Despite interpatient variability in plasma cell viability, profound LC suppression was generally achieved with less cytotoxicity than BZ treatment (Fig. 5C; Supplementary Fig. 3A). Because t(11;14) has been associated with reduced hematologic and organ responses to BZ-based therapy in AL amyloidosis (Bochtler et al., 2015), we examined whether cytogenetic status influenced treatment response. While plasma cell sensitivity to BZ varied significantly with t(11;14) status, responses to Sec61 inhibition were unaffected (Supplementary Fig. 3B). Moreover, unlike BZ, which induced marked toxicity in matched, autologous CD138⁻ bone marrow cells, Sec61 inhibition largely preserved the viability of non-plasma cell populations (Fig. 5D). Together, these findings demonstrate that A317-driven Sec61 inhibition robustly suppresses pathogenic LC production across genetically and clinically diverse AL patient samples while sparing non-producing bone marrow cells.

**Figure 5.**
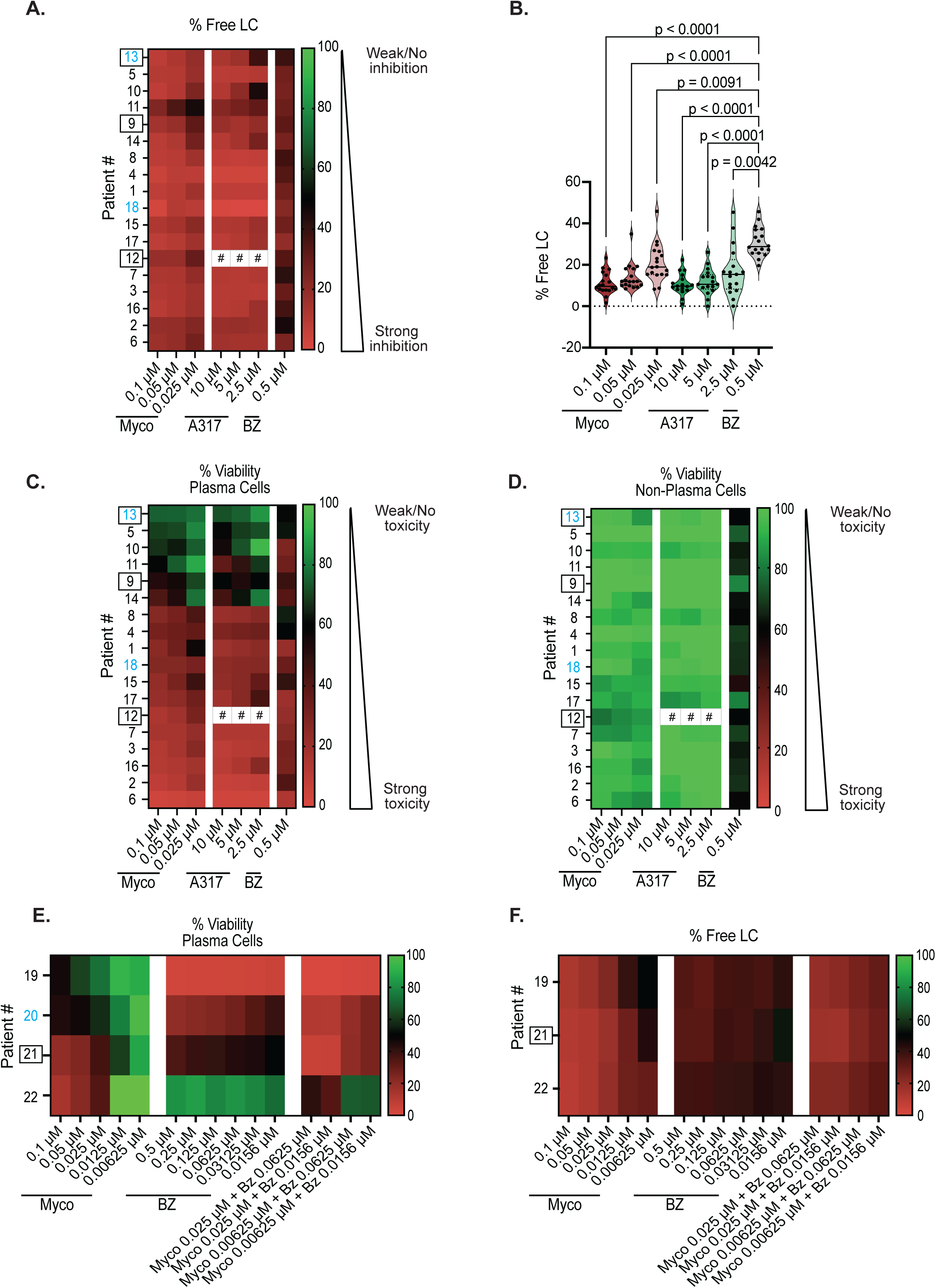
Selective Sec61 inhibition prevents pathogenic LC biogenesis in AL. **(A)** Effects of mycolactone (Myco), A317, and bortezomib (BZ) on free LC secretion by CD138⁺ bone marrow plasma cells from 18 patients with AL amyloidosis. Results are expressed relative to vehicle controls. Blue: patients with κ-LC isotype; black: patients with λ-LC isotype; boxed: previously treated patients; unboxed: treatment-naïve patients. #: not determined, due to an insufficient number of primary cells from the corresponding patient sample. **(B)** Violin-plot representation of the data in (A); each dot represents one patient. Repeated-measures mixed-effects statistical test with Dunnett’s multiple-comparisons test. **(C–D)** Effects of mycolactone (Myco), A317, and BZ on the viability of matched CD138⁺ plasma cells (C) and CD138⁻ non-plasma bone marrow cells (D). **(E–F)** Effects of mycolactone (Myco), BZ, and their combinations on the viability **(E)** and free LC secretion **(F)** of primary CD138⁺ plasma cells from four additional patients with AL amyloidosis. All values are expressed relative to vehicle controls.

We next investigated the interaction between Sec61 and proteasome inhibition in primary AL plasma cells from four additional patients (Table II; patients 19-22). CD138⁺ cells isolated from bone marrow aspirates were treated *ex vivo* with mycolactone (6.25 nM-0.1 µM) or BZ (15.6 nM-0.5 µM), alone or in combination, and the effects on cell viability and LC secretion were assessed as above (Fig. 5E, F). Mycolactone was used for these combination studies as a prototypical Sec61 blocker, given the limited primary cell availability. The four samples spanned a broad range of sensitivity to BZ, with patient 19 exhibiting marked susceptibility to BZ-induced cell death, patient 22 displaying pronounced resistance, and patients 20 and 21 showing intermediate responses. In samples where BZ alone failed to induce maximal cytotoxicity (patients 20, 21, and 22), the addition of low concentrations of mycolactone consistently enhanced BZ-induced plasma cell death, yielding positive synergy scores (Supplementary Fig. 3C-D). Of note, in patient 22, higher mycolactone concentrations antagonized the lowest concentrations of BZ. Collectively, these findings demonstrate that partial Sec61 inhibition potentiates proteasome blockade in primary AL plasma cells, particularly in samples with reduced sensitivity to BZ, while highlighting the importance of dose optimization to maximize therapeutic synergy.

## DISCUSSION

This study identifies Sec61-mediated ER translocation as a therapeutic vulnerability in AL amyloidosis. Using the prototypic inhibitor mycolactone, we show that pharmacological Sec61 blockade prevents ER entry of amyloidogenic LCs and redirects nascent chains to proteasomal degradation, thereby suppressing production of the amyloid precursor at its source. By imposing an additional proteostatic burden on highly secretory plasma cells, mycolactone also enhanced bortezomib activity *in vitro* and *in vivo*. However, mortality in mice receiving the drug combination revealed the narrow therapeutic window of pan-Sec61 inhibition, arguing against broad disruption of the secretory pathway as a clinically viable approach. These findings prompted us to evaluate the newly developed, PD-1-selective Sec61 inhibitors A317 and A347, which revealed that the extent of Sec61 client selectivity determines the balance between therapeutic activity and toxicity.

Although A317 and A347 both inhibited LC translocation through Sec61, proteomic profiling revealed distinct effects not only across the Sec61 clientome but also across the broader plasma-cell proteome. Importantly, the number of significantly modulated proteins was small relative to the more than 7,000 proteins detected under the conditions tested: 2.0% proteins were altered by A317 and 1.1% by A347. Despite limited overlap among individual regulated proteins, both compounds elicited a common, highly restricted signature of Sec61 perturbation. Each reduced LC abundance and MAN2B2 levels, while increasing the ERAD component HERP, the ER-stress sensor IRE1α, and the dolichol-synthesis enzymes DHDDS and NUS1. This shared response is consistent with impaired ER protein entry, altered glycoprotein processing, and compensatory activation of ER proteostasis pathways. A317 elicited a broader and more pronounced version of this response, including reduced MAN1B1 and IL6R expression and increased XBP1 expression, whereas A347 induced a more restricted signature. Thus, both compounds appear to engage a common Sec61-dependent mechanism but differ in the breadth and magnitude of their direct and secondary effects. This difference translated into distinct therapeutic profiles. A317 maintained broader LC-suppressive activity than A347 while remaining substantially more selective than mycolactone and consistently reduced pathogenic LC secretion in primary AL plasma cells with limited toxicity to matched non-plasma cells from bone marrow. These results indicate that intermediate, rather than maximal, Sec61 selectivity may best balance anti-LC activity and tolerability.

SP sequence emerged as a key determinant of this balance. Although mycolactone uniformly inhibited all tested LC SPs, susceptibility to A317 varied among sequences and was markedly more restricted for A347, which failed to inhibit several λ-chain and all κ-chain SPs tested. A317 therefore provided substantially broader coverage of the major amyloidogenic LC germline families. Based on their reported frequencies (Morgan et al., 2025), we estimate that A317 and A347 would cover approximately 70% and 42% of patients with AL amyloidosis, respectively. These observations indicate that increasing selectivity can restrict patient coverage and suggest that highly selective Sec61 inhibitors may require patient stratification based on LC SP sequence.

Beyond direct LC suppression, intermediate-selective Sec61 inhibition may modulate pathways supporting plasma-cell survival and the inflammatory bone marrow niche. A317 retained activity against several Sec61-dependent receptors implicated in these processes, including PD-1, TNFR1, TNFR2, CD25, and IL-6R, whereas A347 showed little or no activity against most of these targets. Although their contribution to therapeutic efficacy remains to be defined, concurrent attenuation of LC production and disease-relevant signaling may be an advantage of intermediate-selective inhibitors. Conversely, the narrower client spectrum of A347 may improve tolerability, but at the expense of patient coverage and potential activity against these additional targets.

Our combination studies suggest that the interaction between Sec61 and proteasome inhibition may depend on both the degree of Sec61 blockade and the intrinsic BZ sensitivity of plasma cells, although these observations derive from a small number of patient samples and should be interpreted with caution. In samples with residual resistance to BZ, low concentrations of mycolactone consistently enhanced BZ-induced cytotoxicity, whereas higher concentrations provided no additional benefit and were antagonistic in one resistant sample, an isolated observation that will require confirmation in a larger cohort before any general conclusion can be drawn. No synergy was observed in the highly BZ-sensitive sample, likely because BZ alone induced near-maximal cell death. Overall, these preliminary findings suggest that the combination’s efficacy may be dose- and context-dependent, warranting further optimization and validation in a larger patient cohort.

Several limitations should be acknowledged. Although key findings were validated in primary patient-derived samples, the mechanistic analyses relied largely on a single AL cell line. In addition, Sec61 client responses in the native cellular context may also be shaped by cell type- and protein-level features not captured by the SP reporter system, including protein half-life, and the efficiency of post-translational processing or degradation. Finally, further medicinal-chemistry optimization, together with pharmacokinetic, pharmacodynamic, and toxicological studies, will be required to determine whether A317 or related compounds possess the properties needed for clinical development.

In conclusion, selective Sec61 inhibition suppresses production of the pathogenic amyloid precursor and represents a mechanistically novel therapeutic strategy for AL amyloidosis. Our findings further show that Sec61 client selectivity is a tunable pharmacological property that governs therapeutic breadth, proteostatic consequences, and tolerability. More broadly, the data support the feasibility of pharmacologically targeting a disease-relevant secretory protein while limiting perturbation of the wider secretory proteome, a central challenge in developing Sec61-directed therapies.

## MATERIALS AND METHODS

### Reagents

Mycolactone was purified from *M. ulcerans* bacterial pellets (strain 1615), then quantified by spectrophotometry, and stored in ethanol at -20°C protected from light. For *in vivo* experiments, a 4 mM stock was diluted in a NaCl solution (0.9% w/v) immediately before injection in animals. For *in vitro* experiments, a 1,000× working solution was prepared by dilution of the ethanol stock in DMSO and stored at -20°C, then thawed and diluted in culture medium immediately before use. BZ purchased from Alfa Aesar (#J60378) was resuspended in DMSO to yield a 10 mM solution stored at -20°C. BZ was thawed and diluted in culture medium immediately before use. A317 (#HY-139615) and A347 (# HY-139616) were purchased from Medchem Express, resuspended in DMSO to a 10 mM stock solution stored at -20°C.

### Cell lines and cultures

The ALMC-1 cell line (Arendt et al., 2008) was purchased from Merck (cat# SCC430) and cultured in IMDM (Gibco) supplemented with 10% fetal bovine serum (Dominique Dutscher), 1% penicillin/streptomycin (Gibco), 1% Sodium pyruvate (Gibco) and 2ng/ml IL-6 (Bio-Techne) at 37°C and 5% CO_2._ HEK293 cells were purchased from The European Collection of Authenticated Cell Cultures (ECACC, cat# 85120602). HEK293 cells were transduced using Lipofectamine 2000 (Thermo Fisher) with the plasmid pCMV3-IGLV6-57-opsin, generated in our lab. A stable cell line was generated by antibiotic selection (hygromycin, 100 µg/mL), called HEK293-LC-opsin. Flp-In™ T-REx™ 293 Cell Line was purchased from ThermoFisher (cat# R78007) and transduced using Lipofectamine 2000 (Thermo Fisher) with the plasmid pTwist-IGLV6-57-nLuc (NanoLuc® Luciferase, Promega) (plasmid and construction made by Twist Bioscience) (LC-nLuc). A stable cell line was generated by antibiotic selection (puromycin, 1 mg/ml). The stable cell line was subsequently co-transduced using Lipofectamine 2000 (Thermo Fisher) with pcDNA5-Sec61A1 WT or R66G plasmids (Itskanov et al., 2023) and pOG44 Flp-Recombinase Expression Vector (ThermoFisher). Stable cell lines were generated by antibiotic selection (hygromycin, 100 µg/mL). To generate FL(IGLV6-57) and SP(IGLV6-57)-expressing cell lines, Flp-In™ T-REx™ 293 Cell Line was co-transduced as described above using the pOG44 and the pcDNA5 vector containing either the full length IGLV6-57 gene or the SP sequence, respectively. To generate cell lines expressing the panel of amyloidogenic κ- and λ-LC signal peptides (SPs), as well as the panel of SPs derived from plasma cell- and inflammatory niche–relevant proteins, Flp-In™ T-REx™ 293 cells were co-transfected as described above with pOG44 and pcDNA5 vectors encoding the respective SPs. Protein sequences were obtained from UniProt: P01721 (IGLV6-57); P01703 (IGLV1-40); P01699 (IGLV1-44); P01704 (IGLV2-14); P01715 (IGLV3-01); P01594 (IGKV1-33); P01597(IGKV1-39); P06312 (IGKV4-01). SP sequences were verified using SignalP 6.0 (DTU Health Tech). At the C-terminus of each identified SP, six amino acids from the corresponding native protein were added upstream of the sequence encoding nLuc. The above cell lines were cultured in DMEM (Gibco) supplemented with 10% fetal bovine serum (Dominique Dutcher), 1% penicillin/streptomycin (Gibco), 1% Sodium pyruvate (Gibco). All cell lines were routinely tested for mycoplasma.

### NanoLuc® Luciferase secretion assay

Test compounds and controls were prepared in white-bottom 96- or 384-well plates (Greiner). An Echo® 555 Liquid Handler (Beckman) was used to dispense compounds. On day 0, cells were seeded in the compounds-containing plates at a density of 8,000 cells/well (96 well) or 2,000 cells/well (384 wells). After 24h of incubation at 37°C with 5% CO_2_, the plates were briefly centrifuged and LC-nLuc secretion in the supernatant was assayed through Nano-Glo® Luciferase Assay System (Promega) following manufacturer’s instruction. Cell viability was assessed through CellTiter-Glo® assay (Promega). Luminescence signal was read using the FLUOstar OPTIMA (BMG Labtech) plate reader or the Tecan Spark plate reader.

### ALMC-1 cells secretion and viability assay

Test compounds and controls were dispensed into white-bottom 96-well plates (Greiner) using an Echo® 555 Liquid Handler (Beckman Coulter), in triplicate. On day 0, ALMC-1 cells were seeded into the compound-containing plates at a density of 50,000 cells/well and incubated for 24 h at 37°C with 5% CO₂. Plates were then centrifuged, and supernatants were collected for quantification of λ LC secretion by ELISA. Cell viability was assessed in parallel using the CellTiter-Glo® assay.

### Detection of secreted immunoglobulin LC by ELISA

Nunc Maxisorp plates (ThermoFisher) were coated with anti-human lambda LC antibody (Bethyl #A80-116A) or anti-Human Lambda Free Light Chain (Bethyl #A80-127) diluted 1/500 in coating buffer (Biolegend) at 4°C overnight. ELISA plates were then washed in 1X Tris Buffered Saline pH 8 (TBS, Fisher)-0.05% Tween 20, 3X and blocked in blocking buffer (1% Bovine Serum Albumin, Interchim in 1X TBS pH 8) for 1h at RT. Cellular supernatants were diluted in blocking buffer. A standard curve was prepared starting from the purified Human Fab/Lambda IgG fragment (Bethyl #P80-116) and purified immunoglobulin Lambda Light Chain (free) from human (Sigma #L0665). Supernatants or standards were added to the ELISA plates and incubated for 1h at RT. After 3 washes, the anti-Human Lambda LC Antibody Biotinylated (Bethyl #A80-116B) diluted 1/100,000 in dilution buffer (blocking buffer with 0.05% Tween 20) was incubated for 1h at RT. Following 3 washes, the streptavidin-HRP (Biolegend) diluted 1/2,000 or 1/3,000 in dilution buffer was incubated for 30 minutes at RT. After 3 washes, TMB substrate (Biolegend) was added in each well. The plates were incubated for 15 minutes at RT, and the reaction was stopped with 2N sulphuric acid solution (VWR). The absorbance of the colorimetric signal was measured at 450nm using the FLUOstar OPTIMA (BMG Labtech) plate reader. Free Kappa Light Chains were detected using a kit (Biovendor #RD194088100R) following manufacturing recommendation.

### Western blot analysis

Cells were treated for 6h with DMSO or compounds +/- proteasome inhibitor MG132 (10µM, Sigma), collected, washed with DPBS and pelleted. To lyse the cells, the following buffer was used: Tris-HCl pH 7.4 20 mM, NaCl 150 mM, n-dodecyl-{beta}-D-maltoside 0.2%, EGTA 1 mM, NaF 50 mM, MgCl2 1 mM, and protease inhibitors cocktail (Sigma) 1X. Proteins were solubilized for 10 minutes on ice and after 10 minutes centrifugation at 16,000 g at 4°C, total amounts of proteins were quantified by NanoDrop Lite (ThermoScientific). 16 mg of total proteins/sample were used for the EndoH treatment (NEB kit #P0702S) following manufacturer’s instructions. Proteins were separated by SDS-PAGE using 4-12% NuPAGE Bis-Tris gels (Invitrogen) and transferred onto nitrocellulose membranes (iBlot 3® gel transfer Stacks Nitrocellulose system from Invitrogen). Immune blotting was carried out with the primary antibodies anti-LC (Bethyl #P80-116) and anti-tubulin (CST #2148S), both diluted 1/1,000 in 1% fat-free milk in PBS-0,1% Tween. After washing, the membranes were incubated with HRP-conjugated anti-rabbit or -goat IgG secondary antibodies (Santa Cruz #sc-2357 and #sc-2922, respectively) at a concentration of 1/5,000 in 1% fat-free milk in PBS-0,1% Tween. Detection of proteins was performed using the SuperSignal^TM^ West Femto (ThermoScientific), and images were acquired on ImageQuant 800 (Amersham). Band intensities were quantified by densitometry in Fiji/ImageJ. The percentage of glycosylated LC was calculated as the intensity of the glycosylated LC band divided by the sum of glycosylated and non-glycosylated LC bands ×100. Total LC was calculated as the sum of glycosylated and non-glycosylated LC band intensities, normalized to α-tubulin, and expressed relative to DMSO.

### Proteomic analysis

For each condition, 10^7^ cells were seeded in 75 cm2 flasks (TPP) in 17 mL of medium. Each condition was performed in five technical replicates. The cells were incubated with DMSO or the compounds for 24h at the concentration corresponding to 50% inhibition of LC secretion, as determined by ELISA (A317 0.3 μM or A347 1.5 μM) at 37°C in 5% CO_2_. Cells were then harvested, lysed and the protein content underwent the same enzymatic digestion protocol as described in the publication by Morel et al. (2018). Peptide solutions were desalted using the AssayMAP Bravo (Agilent) with C18 cartridges (Agilent Technologies, 5 μL bead volume) and eluted with ACN 80 %, FA 0.1 %. Finally, the peptide solutions were speed-vac dried and resuspended in acetonitrile (ACN) 2%, FA 0.1% buffer before injection in a nanochromatographic system (Vanquish Neo - ThermoFisher Scientific) coupled online to an Orbitrap Eclipse tribrid mass spectrometer (ThermoFisher Scientific) equipped with a FAIMS Pro Duo. For each samples, 1 µg of peptides were loaded into a C18 column (EASY-Spray^TM^ - ES903 - ThermoFisher Scientific: 50cm x 75 µm ID, 2.0 µm particles, 100 Å pore size,) after an equilibration step in 100 % solvent A (H2O, 0.1% FA). Peptides were eluted with a multi-step gradient from 5 to 25% buffer B (ACN 80% / FA 0.1%) in 95 min, 25 to 40% buffer B in 15 min and 40 to 95% Buffer B in 10 min at a flow rate of 250 nL/min for up to 130 min. Column temperature was set to 50°C and FAIMS CV set to -48. The Data-Independent Acquisition (DIA) method consisted in a succession of one MS scan (from 350 to 1250 m/z) at a resolution of 60,000 and 40 MS/MS scans of 1 m/z overlapping windows (isolation window = 20 m/z) from 400 to 1200 m/z at 30,000 resolutions. The AGC (Automatic Gain Control) target and maximum injection time for MS and MS/MS scans were set to 4.0E5, 50 ms and 5.0E5, 54 ms respectively. The normalized collision energy was set to 30 for HCD fragmentation. The mass spectrometry proteomics data have been deposited to the ProteomeXchange Consortium via the PRIDE partner repository (Perez-Riverol et al., 2025) with the dataset identifier PXD083851.

### Proteome analysis and statistics

Spectronaut 20.1.250624.9244917.0.221202.55965 (Quasar) (Biognosys AG) was used for DIA-MS data analyses with UniProt homo sapiens database (downloaded on 29/11/2024). The data extraction was performed using the default BGS Factory Settings. Briefly, for identification, both precursor and protein FDR were controlled at 1%. For quantification, Qvalue was used for precursor filtering, and no imputation strategy was used; peptides were grouped based on stripped sequences. Cross Run Normalization was enabled. Statistical analysis compared protein intensities across conditions. Protein quantitative values were measured using Spectronaut (PG.Quantity). To identify proteins that were more abundant in one condition than in another, the quantified values from both conditions were compared. Only proteins with at least four quantitative values in one of the two conditions compared were retained for further statistical analysis to ensure a minimum of replicability. The remaining values were then log2-transformed and normalized by median centering within each condition. Quantitative values associated with at least one peptide were kept for further statistics. Proteins without any quantitative value in one of both conditions have been considered as proteins quantitatively present in one condition and absent in the other. They have, therefore, been set aside and considered differentially abundant proteins. Next, missing values of the remaining proteins were imputed using the impute.mle function of the R package imp4p (Gianetto et al., 2020). Statistical testing was conducted using limma t-tests thanks to the R package limma (Ritchie et al., 2015). The FDR control was performed using an adaptive Benjamini-Hochberg procedure on the resulting p-values thanks to the function adjust.p of R package cp4p using the robust method to estimate the proportion of true null hypotheses among the set of statistical tests (Giai Gianetto et al., 2016). The thresholds to consider a protein differentially abundant are set at a fold-change (FC) of 1.5 (log₂(FC) ≥ 0.58 or ≤ -0.58) and a False Discovery Rate (FDR) of 1%.

### Patients

Clinical records and bone marrow aspirates were obtained from 22 patients with AL amyloidosis referred to the Italian Amyloid Center or to the Department of Hematology – Fondazione Istituto di Ricovero e Cura a Carattere Scientifico Policlinico San Matteo (IRCCS), Pavia. In accordance with the Declaration of Helsinki, all patients gave their written informed consent for the use of their clinical data and biological samples for research purposes, in agreement with the Institutional Review Board guidelines (CET-L6 Protoc. N. 0067361/24). Positive- and negative-CD138 bone marrow plasma cell populations were isolated through immunomagnetic cell sorting using CD138 microbeads, MS columns and an OctoMACS separation system (all from Miltenyi Biotec). CD138^-^ cell populations were isolated after the first round of isolation whereas positive fractions were selected after two sequential rounds of isolation. The purity of cell populations was assessed by flow cytometry after incubation with the anti-CD138-BV421 antibody (Becton Dickinson Biosciences, BD) for 15 minutes, fixation with the BD FACS Lysing solution 10X (Becton Dickinson Biosciences, BD) for 10 minutes and subsequent acquisition on a BD FACSCanto II cytometer. Data analysis was conducted with Flow Jo software (Becton Dickinson Biosciences, BD) (Table 1). CD138^+^ and CD138^-^ populations were seeded at 3 x 10^4^ cells per well in 96-well plates with four technical replicates. After treatment, the plates were centrifuged and supernatant was harvested for secretion studies. Cell viability was assessed through CellTiter-Glo® Luminescent Cell Viability Assay. Luminescence was measured using the Tecan Infinite F200 plate reader. Relative metabolic activity of drug-treated cells was compared to DMSO-negative control (set as 100%).

### Mouse studies

NOD. Cg-Prkdcscid Il2rgtm1Wjl (NSG, stock number: 005557) mice were purchased from the Jackson Laboratory and were used between 8 and 13 weeks of age. All mice were housed at animal facilities of the Institut Pasteur under specific pathogen-free conditions with food and water *ad libitum*. Mouse experiments were validated by CETEA Ethics Committee CEEA89 (Institut Pasteur, Paris, France, DAP240017) and received the approval of the French Ministry of Higher Education and Research. They were performed in compliance with national guidelines and regulations. 2 × 10⁶ human ALMC-1 cells, resuspended in 100 μl of phenol red-free Corning® Matrigel® Matrix (Corning, #356237), were subcutaneously injected into the flank of each mouse under isoflurane anesthesia. Blood microsamples were collected from the tail vein every 3.5 days into EDTA-containing tubes. Plasma was obtained by centrifugation and used to quantify circulating free light chain concentrations by ELISA. Mice received intraperitoneal injections of mycolactone (Myco; 0.3 mg/kg) and/or bortezomib (BZ; 0.5 mg/kg) in 200 μl sterile 0.9% NaCl, or vehicle alone, twice weekly. Graft growth was monitored by unblinded caliper measurements, and volume was expressed in mm³. Animals were euthanized when plasma LC reached a predefined plateau based on prior experiments, or earlier if signs of pain or distress were observed, in accordance with institutional ethical guidelines.

### Histology

Cell mass, heart, pancreas, liver and kidney were fixed in formalin at room temperature for 24– 48 h, transferred to 70% ethanol and embedded in paraffin. Paraffin sections were cut at 3-µm thickness with a microtome and stained with hematoxylin and eosin (H&E). Slides were scanned using the AxioScan Z1 (Zeiss) system and images were analyzed with the Zen v.2.6 software.

### Synergy scores

Drug synergy between mycolactone and bortezomib was assessed using the Bliss independence model or the highest single agent (HSA) model, calculated in Microsoft Excel. Patients-derived BM plasma cells or ALMC-1 cells were treated with each drug individually or in combination across a matrix of concentrations. Cell viability was measured by CellTiter-Glo® Luminescent Cell Viability Assay after 24h.

### Statistical analyses

Other statistical treatments and graphical representations were performed with the Prism software (v 11.0.2, GraphPad Software, San Diego, CA) and values of P ≤ 0.05 were considered significant. Detailed information on the statistical test used and number of replicates is provided in figure legends.

## ACKNOWLEDGEMENTS

We thank Dr. Eunyong Park, UC Berkeley for kindly providing the pcDNA5-Sec61A1 WT and R66G plasmids. We thank staff of the Amyloidosis Research and Treatment Center and Diagnostic facilities of the Fondazione IRCCS Policlinico San Matteo, Pavia, for clinical evaluations. We thank the Chemogenomic and Biological Screening Platform (PF-CCB) and the Central Animal Facility at Institut Pasteur for their technical assistance. This work was supported by Argobio Studio, Enodia Therapeutics (CD’s team) and by grants from the CARIPLO Foundation (#2023-1731) to AN, and from CARIPLO and Telethon Foundation (#5119) to MN. A CC-BY public copyright license has been applied by the authors to the present document and will be applied to all subsequent versions up to the Author Accepted Manuscript arising from this submission, in accordance with the grant’s open access conditions.

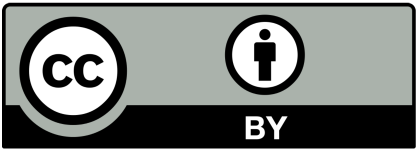

---

**Supplementary Figure 1.**
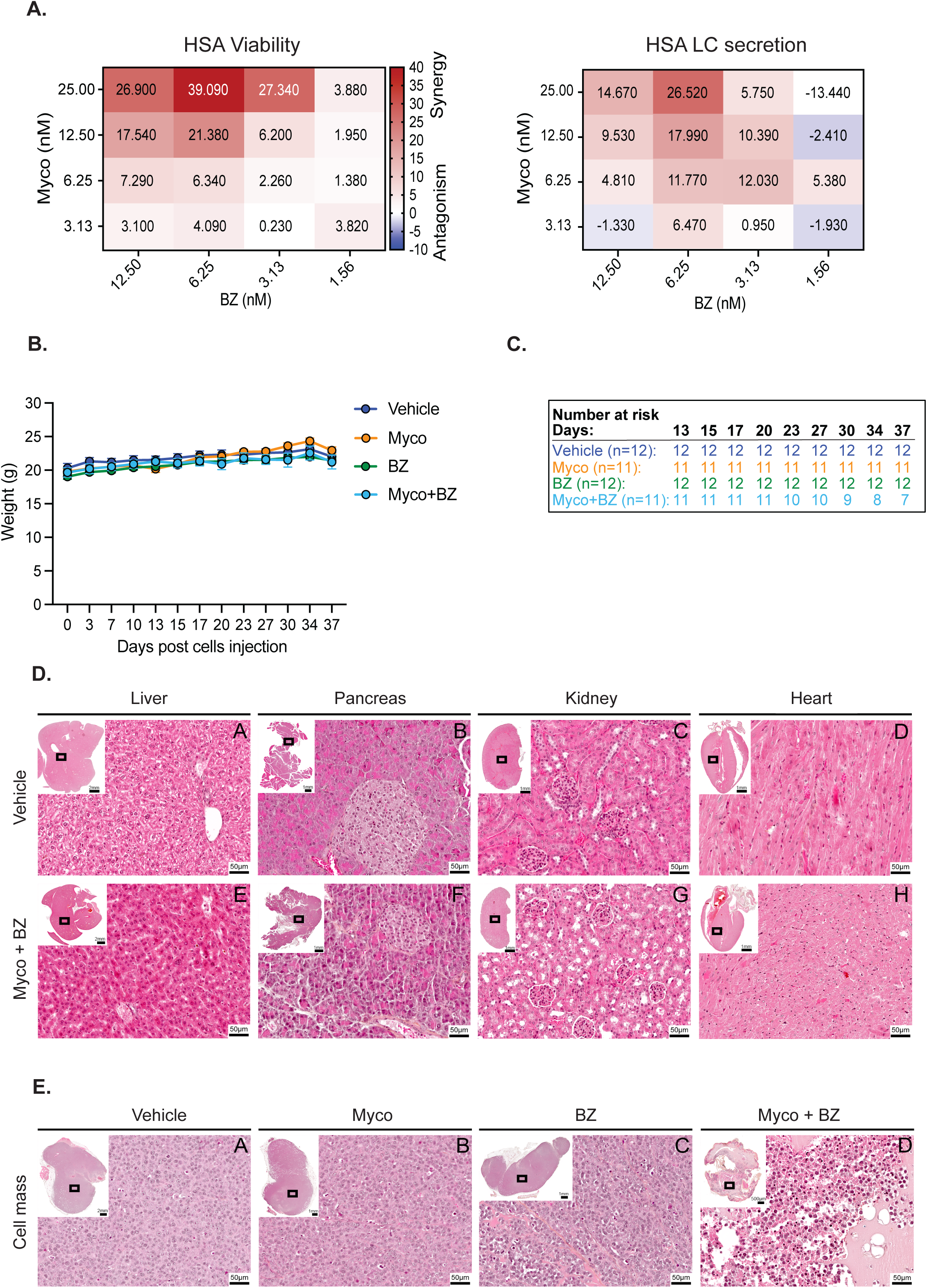
**(A)** HSA synergy scores for the effects of Myco-BZ combinations on ALMC-1 cell viability and free LC secretion. Positive scores indicate synergy. **(B)** Mouse body weight over time, measured at the indicated time points throughout the experiment. Data are shown as mean ± SEM. **(C)** Number-at-risk table for each treatment group, showing the number of mice remaining at each indicated time point throughout the study. **(D)** Representative liver, pancreas, kidney, and heart sections from vehicle- or Myco + BZ-treated mice; insets show the whole organ section, and boxed areas are shown at higher magnification. One organ per mouse is shown per condition. **(E)** Representative tumor cell mass sections from the indicated treatment groups; insets show the whole tumor section, and boxed areas are shown at higher magnification. Scale bars are as indicated.

**Supplementary Figure 2.**
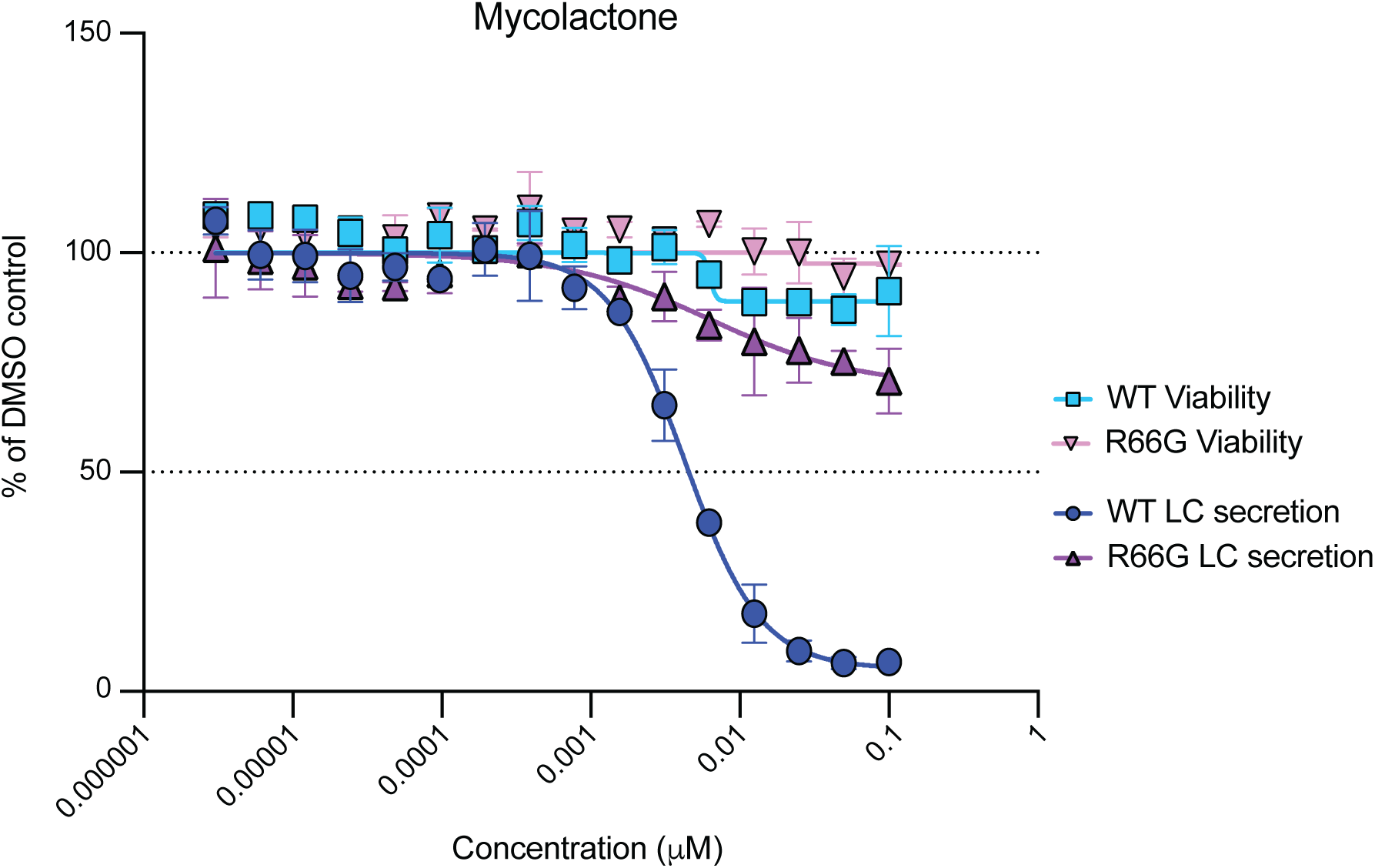
Flp-In T-REx 293 LC–NanoLuc cells expressing wild-type (WT) or R66G Sec61 were treated for 24 h with the indicated inhibitor concentrations. Dose-response curves are expressed relative to vehicle controls. Data are mean ± SD from two independent experiments.

**Supplementary Figure 3.**
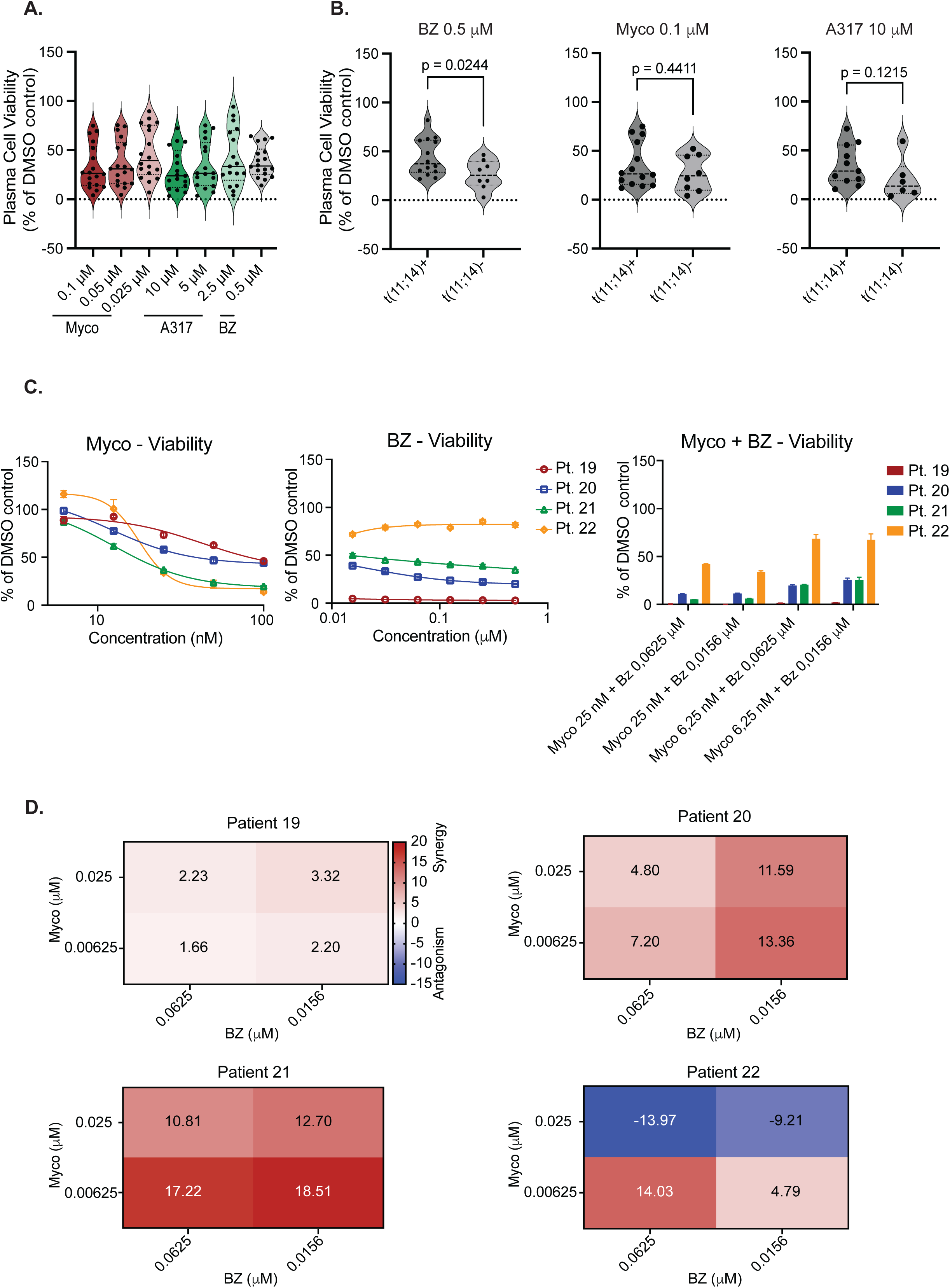
**(A)** AL patients-derived plasma cells show variable sensitivity to Sec61 inhibitors or BZ in terms of cell viability. The data shown in Fig. 4C are presented as a violin plot, with each dot representing one patient. **(B)** BZ, but not Sec61 inhibitors, decreases Plasma Cells viability in a t(11;14)-dependent manner. Data shown in Fig. 4C were grouped according to t(11;14) status and are presented as a violin plot, with each dot representing one patient. Both BZ groups passed the Shapiro-Wilk normality test, and statistical significance was assessed using Welch’s t-test. Differences between t(11;14)⁺ and t(11;14)⁻ samples at 0.1 µM Myco and 10 µM A317 were assessed using a two-tailed exact Mann-Whitney test, given non-normal data distribution; neither comparison reached statistical significance. **(C-D)** In a subset of patients, Mycolactone synergizes with BZ in reducing the viability of AL patients-derived plasma cells. Data shown in Fig. 5E were used to fit dose-response curves (**C**) or calculate Bliss Synergy scores, which are presented as a heatmap (**D**).

